# Vertical and horizontal transmission of cell fusing agent virus in *Aedes aegypti*

**DOI:** 10.1101/2022.05.26.493619

**Authors:** Rhiannon A. E. Logan, Shannon Quek, Joseph N. Muthoni, Anneliese von Eicken, Laura E. Brettell, Enyia R. Anderson, Marcus E.N. Villena, Shivanand Hegde, Grace T. Patterson, Eva Heinz, Grant L. Hughes, Edward I. Patterson

## Abstract

Cell fusing agent virus (CFAV) is an insect specific flavivirus (ISF) found in field and laboratory populations of *Aedes aegypti*. ISFs have recently demonstrated the ability to block the transmission of arboviruses such as dengue, West Nile and Zika viruses. It is thought that vertical transmission is the main route for ISF infections. This has been observed with CFAV, but there is evidence of horizontal and venereal transmission in other ISFs. Understanding the route of transmission can inform strategies to spread ISFs to wild vector populations as a method of controlling pathogenic arboviruses. We crossed individually reared male and female mosquitoes from both a naturally occurring CFAV-positive *Ae. aegypti* colony and its negative counterpart to provide information on maternal, paternal, and horizontal transmission. RT-PCR was used to detect CFAV in individual female mosquito pupal exuviae and was 89% sensitive, but only 41% in male mosquito pupal exuviae. This is a possible way to screen individuals for infection without destroying the adults. Female-to-male horizontal transmission was not observed during this study, however there was a 31% transmission rate from mating pairs of CFAV-positive males to negative female mosquitoes. Maternal vertical transmission was observed with a filial infection rate of 93%. The rate of paternal transmission was 85% when the female remained negative, 61% when the female acquired CFAV horizontally, and 76% overall. Maternal and paternal transmission of CFAV could allow the introduction of this virus into wild *Ae. aegypti* populations through male or female mosquito releases, and thus provides a potential strategy for ISF-derived arbovirus control.

## Introduction

The genus *Flavivirus* contains many arboviruses of medical and veterinary importance such as dengue, West Nile and Zika viruses, as well as insect-specific flaviviruses (ISFs) that are only known to infect insect hosts. ISFs are known to be present in a range of mosquito species and we are becoming increasingly aware of their functions and uses because of recent reports showing their ability to block arbovirus transmission in mosquitoes and their use as chimeric vaccines (Harrison et al. 2021; Baidaliuk et al. 2019; Hobson-Peters et al. 2019; Romo et al. 2018; Hall-Mendelin et al. 2016; Goenaga et al. 2015; Kenney et al. 2014; Hobson-Peters et al. 2013).

Whilst both arboviruses and ISFs belong to the *Flavivirus* genus, transmission is markedly different. Arboviruses are dual-host flaviviruses transmitted through the blood-feeding of arthropods on viremic vertebrate hosts. After acquisition of the virus from the host, only a small fraction of subsequent transmission is vertical, with female mosquitoes passing the virus to their offspring after feeding on an infected host (Phumee et al. 2019; Tesh et al. 2016; Thangamani et al. 2016; Baqar et al. 1993; Rosen et al. 1983). Contrary to this, vertical transmission is thought to be the main route of ISF transmission because the viruses do not replicate in vertebrate hosts.

A small number of studies have observed vertical transmission of ISFs in naturally or experimentally infected mosquito colonies (Peinado et al. 2022; McClean et al. 2021; Contreras-Gutierrez et al. 2017; Bolling et al. 2012; Saiyasombat et al. 2011; Lutomiah et al. 2007), including for cell fusing agent virus (CFAV), an ISF identified in several *Aedes aegypti* cell culture lines, and in field and laboratory colony mosquitoes (Baidaliuk et al. 2020; Martin et al. 2020; Bolling et al. 2015; Stollar and Thomas, 1975). The observed vertical transmission rate was higher in naturally infected colonies of *Ae. aegypti* with CFAV compared to experimentally infected colonies, and similar results were seen in *Culex pipiens* with Culex flavivirus (CxFV) (Contreras-Gutierrez et al. 2017; Saiyasombat et al. 2011). Filial transmission from experimentally infected females ranged from 0-50% for CFAV in *Ae. aegypti*, and 0-22% for CxFV in *Cx. pipiens*. For Kamiti River virus (KRV), an ISF isolated from *Aedes macintoshi*, the filial infection rate from *Ae. aegypti* females following an infectious blood meal was 4% (Lutomiah et al. 2007). The disparate range in these experiments suggest that other forms of transmission may also occur.

Horizontal transmission of ISFs has also been observed in mosquito colonies. KRV was able to infect a high proportion of *Ae. aegypti* larvae when exposed to infected cell culture, while Aedes flavivirus (AeFV) only infected a low proportion of *Ae. aegypti* larvae and adults when feeding on infected cell cultures and sugar meals, respectively (Peinado et al. 2022; Lutomiah et al. 2007). However, the virus was not detected in water used to rear CxFV-infected *Cx. pipiens* larvae and infection was not detected in co-reared, negative larvae (Bolling et al. 2012), suggesting infected individuals did not shed virus into their larval environment. Venereal transmission of CxFV and AeFV was demonstrated in both directions, from male-to-female as well as female-to-male, in experiments that crossed infected mosquito colonies with naïve colonies (Peinado et al. 2022; Bolling et al. 2012). These rates were generally low, except for male-to-female crosses in *Aedes albopictus* which lead to an 18% infection rate (Peinado et al. 2022).

Further knowledge of the transmission and maintenance of ISFs in mosquito populations, especially for mosquito species that are competent vectors where ISFs are of interest for pathogen control approaches, are of high relevance for this group of viruses. CFAV infects an important vector species, *Ae. aegypti*, and its relation to many dangerous flaviviruses may allow it to be used to control arbovirus transmission through superinfection exclusion – blocking subsequent infection of a similar virus – (Baidaliuk et al. 2019) or as a vehicle for paratransgenesis – using a microbe to express transgenes in its host – as has been proposed for other insect-specific viruses (Patterson et al. 2021; Patterson et al. 2020; Gu et al. 2010; Ren et al. 2008; Ward et al. 2001). CFAV has been shown to be maternally transmitted with experimentally infected female mosquitoes (Contreras-Gutierrez et al. 2017), but it is not known if CFAV is paternally or horizontally transmitted. Given the reduced rates of transmission seen in experimental infections, we hypothesized that multiple modes of transmission occur in naturally infected colonies. To assess this, we used a laboratory colony of CFAV-infected *Ae. aegypti* and a known uninfected colony to quantify maternal, paternal, and horizontal transmission of CFAV. Our results provide insights into the transmission routes of CFAV which could be used to inform strategies to spread pathogen-blocking ISFs into mosquito populations.

## Materials and Methods

### Mosquitoes and virus

Established laboratory colonies of *Ae. aegypti* Galveston and Iquitos colonies (kindly provided by Prof. Nikos Vasilakis from the University of Texas Medical Branch) were maintained in a 12-hour light:12-hour dark cycle with a 1-hour dawn and dusk, at 25°C and 75% relative humidity. A previous report identified a persistent infection of CFAV in the *Ae. Aegypti* Galveston colony and no virus infection was detected in the *Ae. aegypti* Iquitos colony (Ma et al. 2021; Bolling et al. 2015). The presence and absence of CFAV in these colonies was confirmed prior to performing the following experiments. The sequence for the CFAV isolate from the Galveston colony is available on GenBank (CFAV-Galveston strain accession no. KJ741267).

### Mosquito rearing to assess vertical and horizontal CFAV transmission

Eggs collected from standard colony maintenance were floated out and larvae were fed ground fish food until reaching pupal stage. Pupae from each colony were sexed and individually placed in 50 mL conical tubes with fresh water. Once emerged, water was removed from the tube and the pupae exuviae were stored at -80 °C, the sex of the adults was confirmed, and individual males were removed from their tube and placed in a tube with an individual female. Mating pairs consisted of the following cohorts: i) 1 Galveston female + 1 Galveston male for CFAV transmission positive control; ii) 1 Iquitos female + 1 Iquitos male for CFAV transmission negative control; iii) 1 Galveston female + 1 Iquitos male for female-to-male horizontal transmission and maternal transmission; or iv) 1 Iquitos female + 1 Galveston male for male-to-female transmission and paternal transmission. Individual mating pairs were provided with 10% sucrose and allowed to mate for three days before the males were removed and stored at -80 °C. Subsequent replicates to confirm the lack of female-to-male transmission involved cohousing mating pairs from cohort iii) 1 Galveston female + 1 Iquitos male for 14 days. Females were presented with a blood meal consisting of 1:1 human red blood cells and plasma via a Hemotek membrane feeding system to stimulate egg laying. After two days, individual blood fed females were transferred to a 30 mL egg laying tube containing water and filter paper. After another two days, females were collected and stored at -80 °C. Egg papers were collected and dried for storage in the insectary until ready to rear offspring. The offspring were reared as normal and emerged adults were collected and individually stored at -80 °C. Adults in the F0 or F1 generation that were deceased prior to collection were not used for detection of CFAV.

### Detection of CFAV by reverse transcription polymerase chain reaction (RT-PCR)

Pupae exuviae or adults were homogenized in a 2 mL Safe-lock microcentrifuge tube with a stainless-steel ball and RNA lysis buffer from the Zymo Quick-RNA Miniprep kit for 5 min at 26 Hz. RNA purification with the Zymo Quick-RNA Miniprep kit was performed per the manufacturer’s protocol.

One-step RT-PCR assays without denaturation were prepared with the Jena Bioscience SCRIPT RT-PCR kit according to the manufacturer’s instructions using CFAV forward and reverse primers as previously described (Weger-Lucarelli et al. 2018). Thermocycler settings were as follows: 1 h at 50 °C, 5 min at 95 °C, 40 cycles of 10 s at 95 °C, 20 s at 60 °C and 2 min at 72 °C, with a final extension of 5 min at 72 °C. An expected amplicon of 367 bp was visualized by gel electrophoresis.

### Statistical Analysis

All statistical analyses were conducted in Microsoft Excel and the R statistical software package (http://www.r-project.org; R Core Team 2020). Binomial 95% confidence intervals (CI) were calculated for sensitivity and transmission efficiencies. Graphics were generated using the package ggplot2 (Wickham 2016).

## Results

### Detection of CFAV in pupal exuviae

Assessment of horizontal and vertical transmission of CFAV requires the confirmation of infected and non-infected mosquitoes prior to the transmission event. This is typically performed by surveying mosquitoes from the colonies to determine a baseline colony infection rate, rather than detection of virus in the individuals involved in the experiment. To assess the experimental individuals directly, pupae exuviae of the parental generation were tested for the presence of CFAV (Figure 1). The pupae exuviae from all CFAV-negative adults were negative, which indicates that the PCR assay has a specificity of 100% in both female and male pupae exuviae. This includes 17/17 females and 47/47 males from the Iquitos colony, and 7/7 females and 1/1 male from the Galveston colony. Only individuals from the Galveston colony were considered for comparison between pupae exuviae and CFAV-positive adults. The resulting sensitivity was 89% (33/37; CI: 74-97%) for females and 41% (11/27; CI: 22-61%) for males, with an overall sensitivity of 69% (44/64; CI: 58-80%) (Table 1). The positive predictive value for CFAV detection in pupae exuviae was 100% for both females and males, and the negative predictive value was 86% for females, 75% for males, and 78% overall. When only considering samples from the Galveston colony, the negative predictive value is 64% for females, 6% for males, and 29% overall.

**Figure 1.**
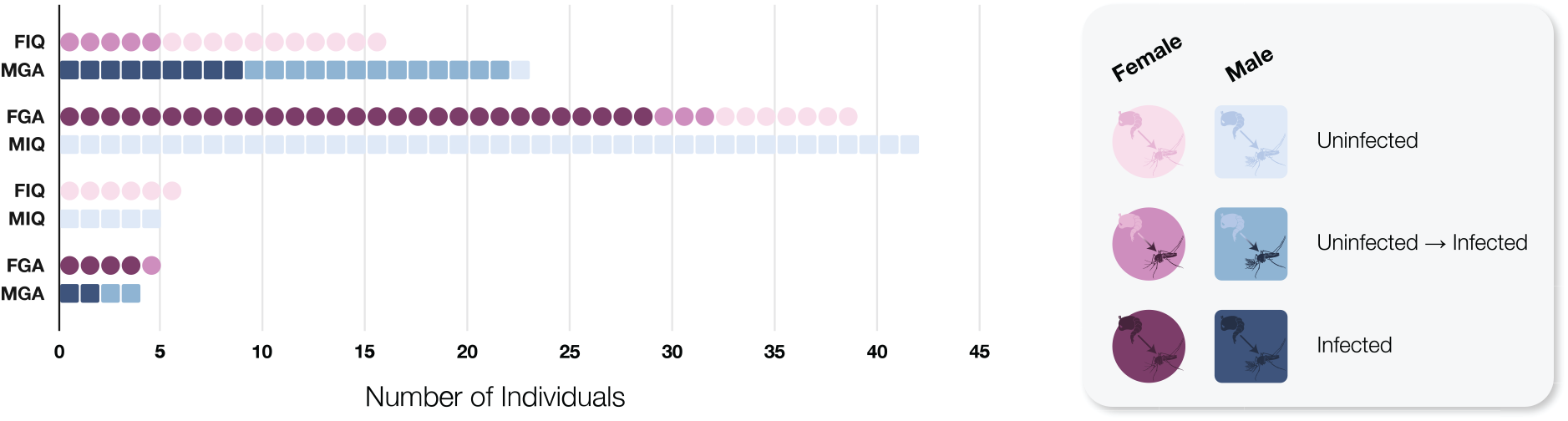
Comparison of CFAV infection status in pupal exuviae versus adult. Red circles indicate the female mosquito and blue squares represent the male in each grouped mating pair. Only mosquitoes that survived to the collection time point were tested, leading to an uneven number of males and females. The shading gradient indicates infection status, where the lightest shade indicates no infection detected in the pupae exuviae and adult, intermediate shade indicates a no infection detected at pupae and positive for infection at adult (pupae result does not agree with adult result), and the darkest shade indicates positive for infection detected at both pupae and adult stages. The combination of each individual mating pair is provided on the Y-axis. The 5 females with negative pupae and positive adult in the FIQ-MGA group are cases of horizontal transmission. FIQ = female Iquitos, MIQ = male Iquitos, FGA = female Galveston, MGA = male Galveston.

**Table 1.**
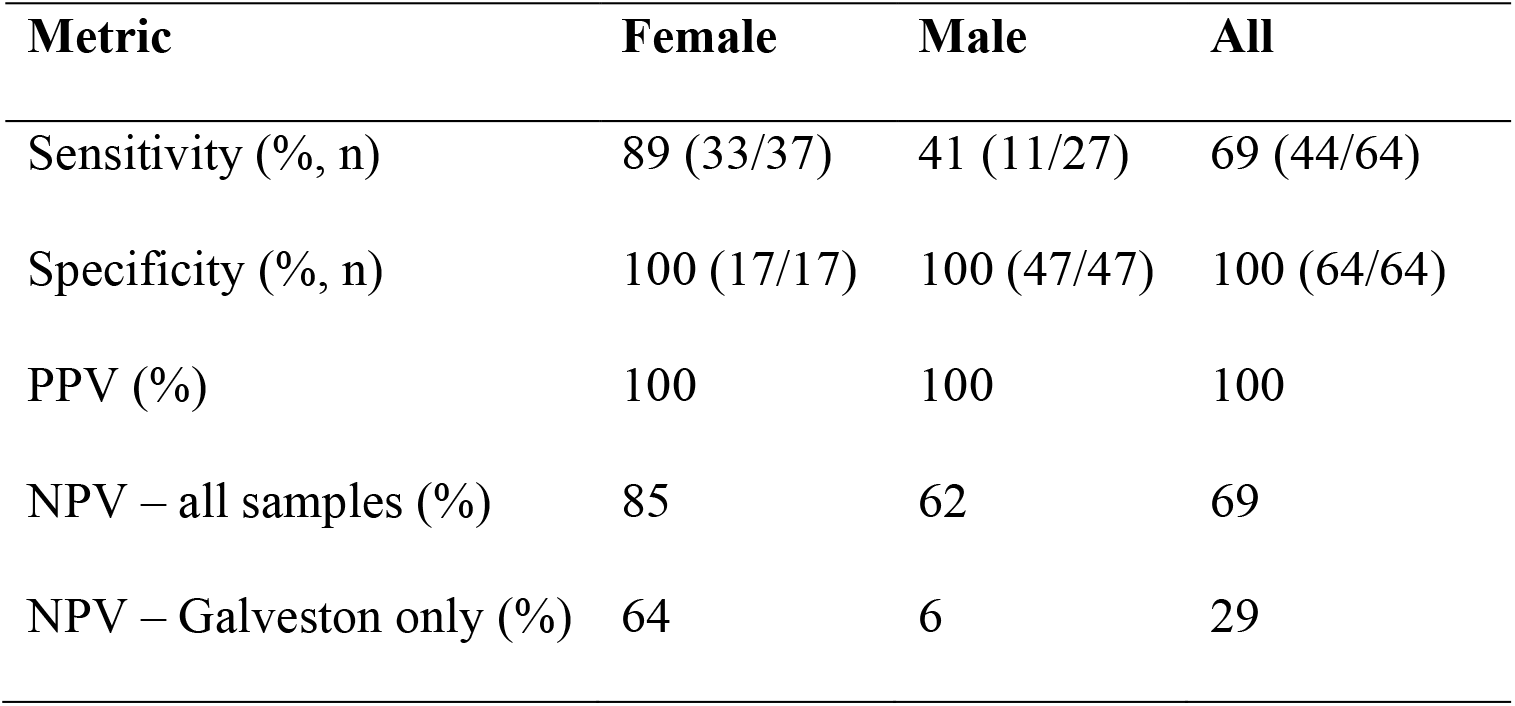
Analysis of detection of CFAV in pupae exuviae versus emerged adult mosquitoes. PPV = positive predictive value, NPV = negative predictive value.

### Horizontal transmission in paired mosquitoes

Adults from the parental generation were grouped into mating pairs and assessed for CFAV infection (Figure 2). All paired adults from the Galveston colony positive control group were positive, and all paired adults from the Iquitos colony negative control group were negative for CFAV. No female-to-male transmission was observed in mating pairs consisting of a Galveston female and an Iquitos male when cohoused for either 3 days (0/14) or 14 days (0/15). All Galveston females were positive, and all Iquitos males were negative for CFAV. Male-to-female transmission was observed at a rate of 31% (5/16; CI: 11-59%) in mating pairs with an Iquitos female and a Galveston male. All Galveston males in these mating pairs were positive for CFAV. The pupae exuviae corresponding to Iquitos female adults where CFAV was detected were all negative.

**Figure 2.**
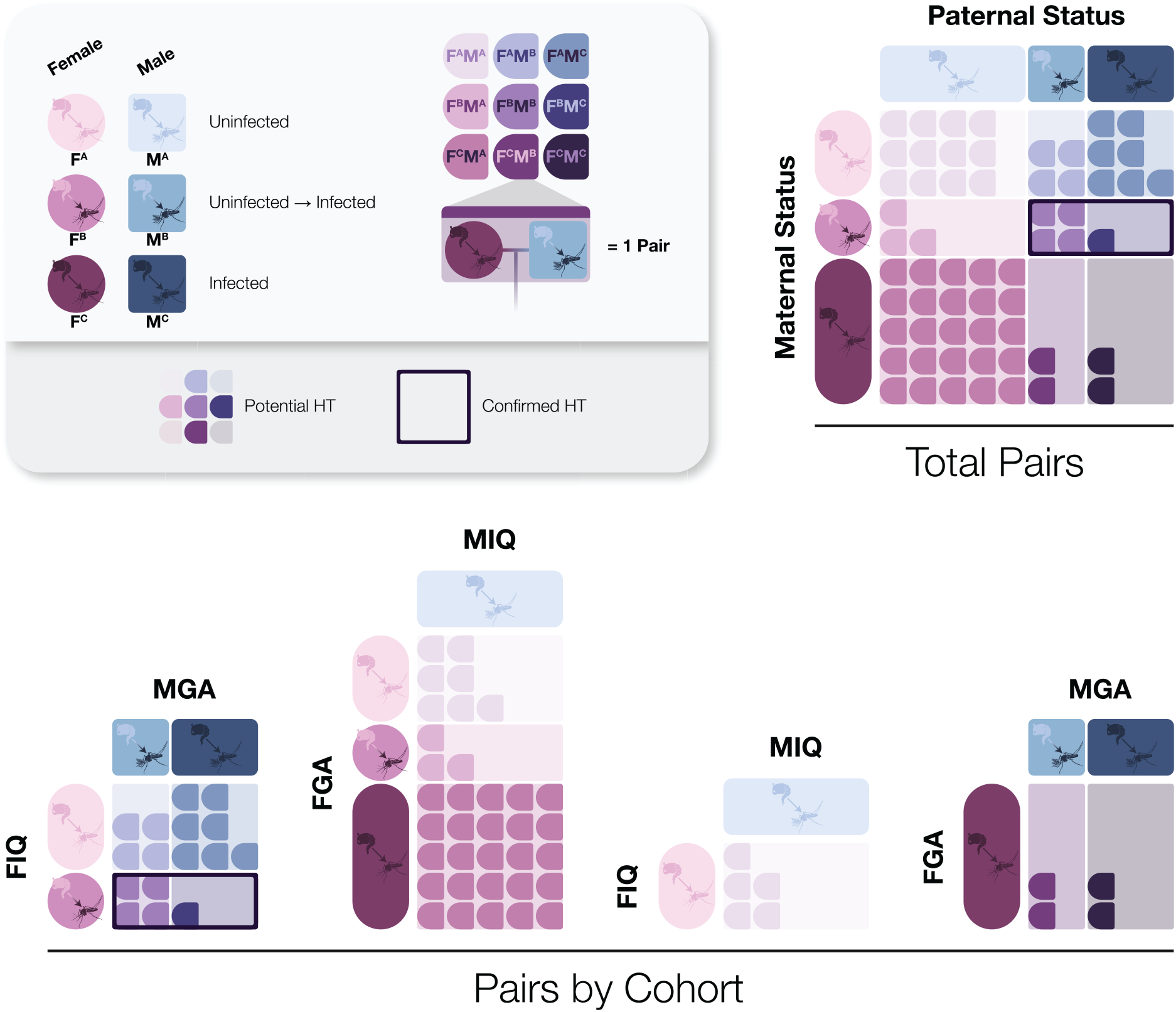
Evidence of horizontal transmission of CFAV between mating pairs. CFAV infection status of F0 mosquitoes in each mating pair. Shading of red circles and blue squares represent the infection status of each adult in the mating pair, with the lightest icons (F^A^ and M^A^) indicating that both mosquitoes in the mating pair tested negative at both the pupa and adult stage, and the darkest icons (F^C^ and M^C^) indicating that both mosquitoes in the mating pair tested positive at both the pupa and adult stage. Mating pairs are assigned a colour on the red-to-blue gradient based on the combined infection status of the female and male (9 potential outcomes). Mosquitoes that tested negative at the pupa stage and positive as an adult (F^B^ or M^B^) are examples of potential horizontal transmission. Samples surrounded by the black border are confirmed cases of horizontal male-to-female transmission. FIQ = female Iquitos, MIQ = male Iquitos, FGA = female Galveston, MGA = male Galveston.

### Vertical transmission from paired mosquitoes

Offspring from all four mating pair groups were assessed for CFAV to determine if vertical transmission was possible through both maternal and paternal routes (Figure 3). Vertical transmission from the Galveston colony control group was 100% (56/56; CI: 94-100%) for offspring from three different mating pairs (Figure 4). Offspring from three Iquitos colony control mating pairs were all negative for CFAV (0/39; CI: 0-9%). Maternal transmission was observed from five mating pairs with a Galveston female and an Iquitos male. The filial infection rate from the five mating pairs ranged from 80-100%, with an overall filial infection rate of 93% (63/68; CI: 84-98%). Paternal transmission was also observed from eight mating pairs with an Iquitos female and a Galveston male, including three mating pairs in which the Iquitos female also became positive. The filial infection rate from mating pairs where the Iquitos female was negative varied from 33-100%, with an overall rate of 85% (56/66; CI: 74-92%). For the three mating pairs with positive Iquitos female adults, the filial infection rate varied from 25-80% with an overall rate of 61% (23/38; CI: 43-76%). The overall filial infection rate from all eight mating pairs with Iquitos female and Galveston male was 76% (79/104; CI: 67-84%).

**Figure 3.**
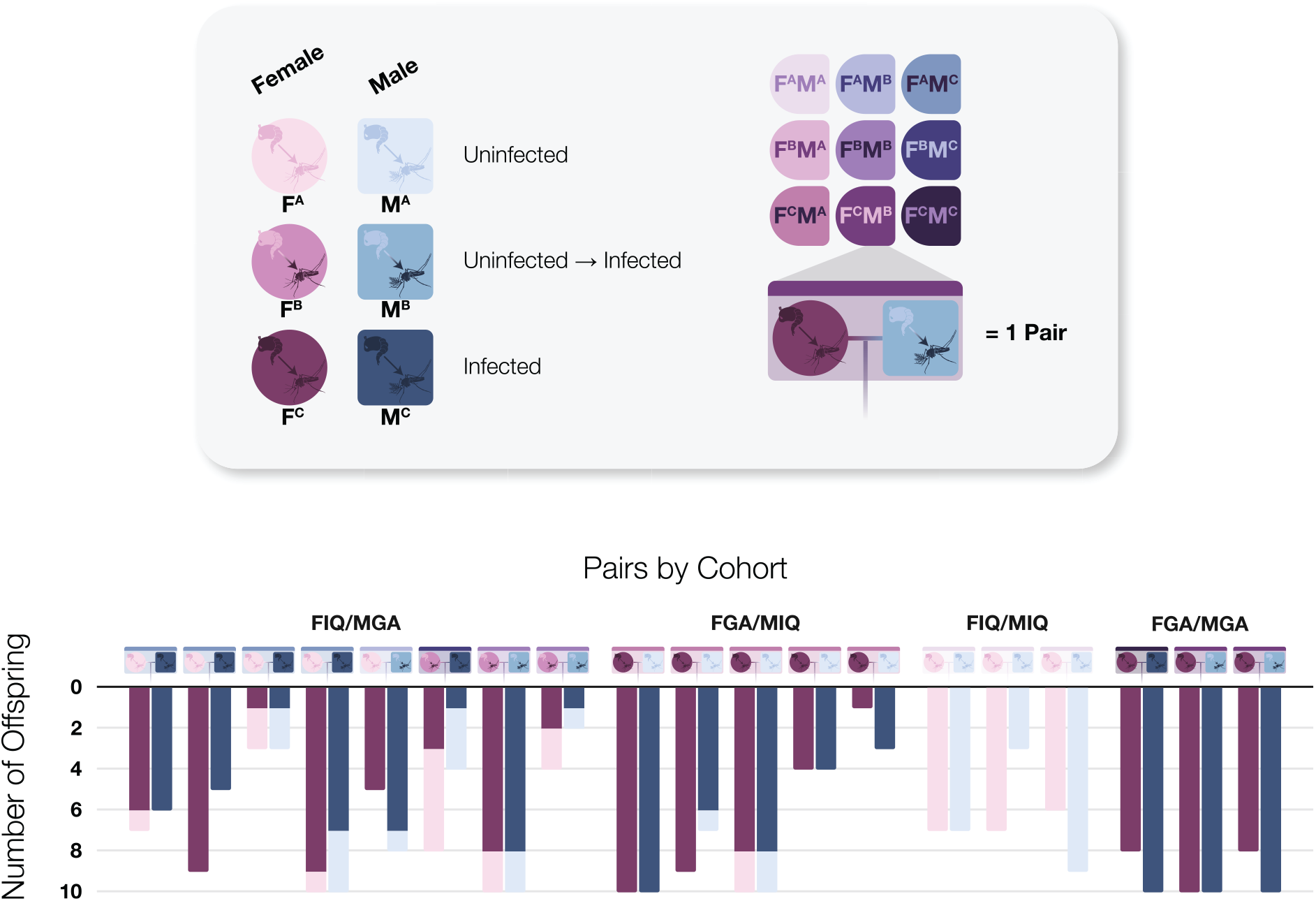
Filial infection of CFAV to assess maternal and paternal transmission. Red circles and blue squares indicate the female and male in each grouped mating pair, respectively. The shading gradient indicates infection status. Shading of red and blue on the bar above the icons represent the infection status of each pupa and adult in the mating pair. Histogram represents the infection status of the adult offspring, with light red or blue representing a negative female or male, respectively, and dark red or blue representing a positive female or male. Pupae exuviae were not examined for offspring. FIQ = female Iquitos, MIQ = male Iquitos, FGA = female Galveston, MGA = male Galveston.

**Figure 4.**
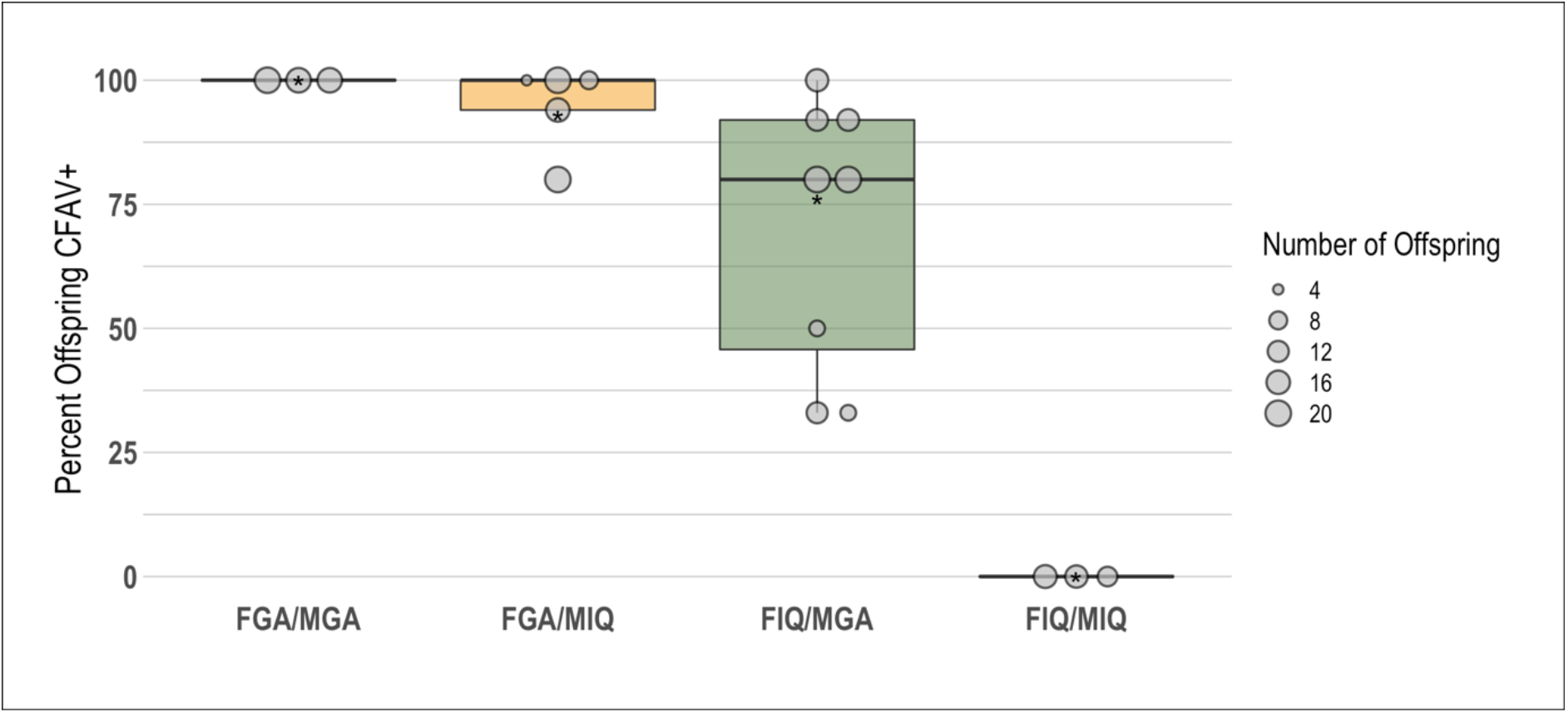
Vertical transmission of CFAV from different mating pair combinations. The size of the grey dots indicates the number of offspring from each mating pair in the group. Asterisk indicates the overall mean of vertical transmission seen from all offspring from the group. FIQ = female Iquitos, MIQ = male Iquitos, FGA = female Galveston, MGA = male Galveston.

## Discussion

There has been a rapid expansion of known members of ISFs and other insect-specific viruses, but little is known about their biology and maintenance in mosquito populations. This is true even for CFAV, an ISF first discovered in 1975 and with global distribution in a major vector species and sustained seasonal infection (Baidaliuk et al. 2020; Jeffries et al. 2020; Martin et al. 2020; Ajamma et al. 2018; Fernandes et al. 2018; Bolling et al. 2015; Yamanaka et al. 2013; Espinoza-Gomez et al. 2011; Cook et al. 2006). The detection of infected larvae or pupae and lack of other known hosts has led to speculation that ISFs are vertically transmitted. Although the results vary by virus and mosquito colony, experimental infections have demonstrated that maternal transmission occurs, as well as venereal transmission and the potential for other modes of horizontal transmission.

Crossing mosquitoes from CFAV-positive and -negative colonies confirmed maternal transmission and revealed paternal and horizontal transmission of CFAV. Maternal transmission of CFAV was first demonstrated by Contreras-Gutierrez et al. (2017). Adult females from a CFAV-negative colony were injected with CFAV, which produced an overall F1 filial infection rate of 28%, and range of 0-50% for individual females. Rearing offspring from the F1 generation increased the overall filial infection rate to 74% in the F2 generation (range of 60-93%), similar to the control rates of 78% to 100% from previous experiments with the Galveston colony (Contreras-Gutierrez et al. 2017) and the current rate of 100% in our Galveston colony. The increase from F1 to F2 infection rates in the experimentally infected colony may be due to the contributions of undetected paternal transmission and chronic infection of CFAV increasing the likelihood of infecting reproductive organs in F1 mosquitoes. Similarly, discrepancies between maternal transmission of 28% compared to 93% in the current experiments may be because the chronic infection of CFAV in Galveston females are more likely to infect reproductive organs compared to injection and 4-day incubation period employed in prior experiments, although ovaries from *Cx. pipiens* were infected with CxFV 4 days post-injection (Saiyasombat et al. 2011). Allowing sufficient time for systemic infection has also been suggested with Anopheles gambiae densovirus (AgDNV), where vertical transmission was observed when parent mosquitoes were infected at the larval stage, but not when females were infected through venereal transmission (Werling et al. 2022; Ren et al. 2008). The high levels of vertical transmission seen in the Galveston colony are also maintained by paternal transmission, which was responsible for an overall filial infection rate of 76% and is the first observation of paternal transmission in ISFs. While paternal transmission has not previously been evaluated in ISFs there are other well-documented examples of paternal transmission, such as for verdadero virus, a partitivirus in mosquitoes (Cross et al. 2020) and rice gall dwarf virus, an aphid-plant reovirus that binds to host sperm to infect offspring (Mao et al. 2019).

Horizontal transmission has also been demonstrated as a viable transmission route for ISFs. Transmission rates from male-to-female adults were 31% for CFAV. This rate is more similar to the 18% male-to-female venereal transmission for AeFV in *Ae. aegypti* than the 2.4% rate observed with CxFV in *Cx. pipiens* (Peinado et al. 2022; Bolling et al. 2012). Neither the current CFAV experiments nor the AeFV experiments excluded other forms of contact transmission, such as sharing sugar meal sources. While transmission through food sharing did not occur with CxFV and feeding on infected sugar meals rarely resulted in AeFV infection (Peinado et al. 2022; Bolling et al. 2012), KRV is known to have high oral infection rates (Lutomiah et al. 2007). No female-to-male transmission was observed, although this occurred at a rate of 5.3% with CxFV, and 2% with AeFV (Peinado et al. 2022; Bolling et al. 2012). Increasing the sample size may reveal some female-to-male transmission, but the rate is likely low.

Our horizontal transmission results were strengthened by testing pupae exuviae to demonstrate the lack of infection before being cohoused and mating with a positive male. Prior studies have not confirmed the infection status of individual mosquitoes before the potential transmission events. Although improvements for testing male pupae exuviae would be desirable, the sensitivity of 89% for female pupae exuviae provides the ability to assess prior infection in mosquitoes by testing pupae exuviae, which will be useful for future experiments. It is unknown why sensitivity differs between female and male pupae exuviae, but a previous study has shown that virus levels have a wider range and are lower titer, on average, in males (Martin et al. 2020).

CFAV may be a useful tool to limit secondary infections with arboviruses in *Ae. aegypti* mosquitoes. Superinfection exclusion has been demonstrated in cells and mosquitoes infected with CFAV, other ISFs and insect-specific viruses. Previous studies showed that initial infection with a field-derived CFAV isolate resulted in reduced dengue virus and Zika virus replication and dissemination (Baidaliuk et al. 2019). However, the lack of knowledge on the transmission of ISFs is a limitation in their potential use for pathogen control (Patterson et al. 2020). As both maternal and paternal transmission have been confirmed, our results offer the potential to establish CFAV infection in wild *Ae. aegypti* populations through the release of infected females or males. Releasing males would be most desirable because they do not contribute to the transmission of arboviruses and CFAV transmission by the male-to-female horizontal route may also improve overall infection levels in the field.

## Acknowledgments

EIP was supported by the Liverpool School of Tropical Medicine Director’s Catalyst Fund award and Brock University Research Initiative Award. JNM was supported by the Wellcome Trust International Masters fellowship (219672/Z/19/Z). GLH was supported by the BBSRC (BB/T001240/1 and BB/W018446/1), the UKRI (20197), the EPSRC (V043811/1), a Royal Society Wolfson Fellowship (RSWF\R1\180013), and the NIHR (NIHR2000907). SH was supported by a Liverpool School of Tropical Medicine Director’s Catalyst Fund award. EH, LEB and GLH were supported by the BBSRC (V011278/1). We would like to thank Prof. Nikos Vasilakis for providing the mosquito colonies used in these experiments.

## References

1. Harrison, J. J., Hobson-Peters, J., Bielefeldt-Ohmann, H., & Hall, R. A. (2021). Chimeric Vaccines Based on Novel Insect-Specific Flaviviruses. Vaccines, 9(11), 1230.

2. Baidaliuk, A., Miot, E. F., Lequime, S., Moltini-Conclois, I., Delaigue, F., Dabo, S., … & Lambrechts, L. (2019). Cell-fusing agent virus reduces arbovirus dissemination in Aedes aegypti mosquitoes in vivo. Journal of virology, 93(18), e00705–19.

3. Hobson-Peters, J., Harrison, J. J., Watterson, D., Hazlewood, J. E., Vet, L. J., Newton, N. D., … & Hall, R. A. (2019). A recombinant platform for flavivirus vaccines and diagnostics using chimeras of a new insect-specific virus. Science translational medicine, 11(522).

4. Romo, H., Kenney, J. L., Blitvich, B. J., & Brault, A. C. (2018). Restriction of Zika virus infection and transmission in Aedes aegypti mediated by an insect-specific flavivirus. Emerging microbes & infections, 7(1), 1–13.

5. Hall-Mendelin, S., McLean, B. J., Bielefeldt-Ohmann, H., Hobson-Peters, J., Hall, R. A., & van den Hurk, A. F. (2016). The insect-specific Palm Creek virus modulates West Nile virus infection in and transmission by Australian mosquitoes. Parasites & Vectors, 9(1), 1–10.

6. Goenaga, S., Kenney, J. L., Duggal, N. K., Delorey, M., Ebel, G. D., Zhang, B., … & Brault, A. C. (2015). Potential for co-infection of a mosquito-specific flavivirus, Nhumirim virus, to block West Nile virus transmission in mosquitoes. Viruses, 7(11), 5801–5812.

7. Kenney, J. L., Solberg, O. D., Langevin, S. A., & Brault, A. C. (2014). Characterization of a novel insect-specific flavivirus from Brazil: potential for inhibition of infection of arthropod cells with medically important flaviviruses. The Journal of general virology, 95(0 12), 2796.

8. Hobson-Peters, J., Yam, A. W. Y., Lu, J. W. F., Setoh, Y. X., May, F. J., Kurucz, N., … & Hall, R. A. (2013). A new insect-specific flavivirus from northern Australia suppresses replication of West Nile virus and Murray Valley encephalitis virus in co-infected mosquito cells. PloS one, 8(2), e56534.

9. Phumee, A., Chompoosri, J., Intayot, P., Boonserm, R., Boonyasuppayakorn, S., Buathong, R., … & Siriyasatien, P. (2019). Vertical transmission of Zika virus in Culex quinquefasciatus Say and Aedes aegypti (L.) mosquitoes. Scientific reports, 9(1), 1–9.

10. Tesh, R. B., Bolling, B. G., & Beaty, B. J. (2016). Role of vertical transmission in mosquito-borne arbovirus maintenance and evolution. Arbovirus: Molecular Biology, Evolution and Control; Vasilakis, N., Gubler, DJ, Eds, 191–217.

11. Thangamani, S., Huang, J., Hart, C. E., Guzman, H., & Tesh, R. B. (2016). Vertical transmission of Zika virus in Aedes aegypti mosquitoes. The American journal of tropical medicine and hygiene, 95(5), 1169.

12. Baqar, S., Hayes, C. G., Murphy, J. R., & Watts, D. M. (1993). Vertical transmission of West Nile virus by Culex and Aedes species mosquitoes. NAVAL MEDICAL RESEARCH UNIT NO 3 CAIRO (EGYPT) DEPT OF MEDICAL ZOOLOGY.

13. Rosen, L., Shroyer, D. A., Tesh, R. B., Freier, J. E., & Lien, J. C. (1983). Transovarial transmission of dengue viruses by mosquitoes: Aedes albopictus and Aedes aegypti. The American journal of tropical medicine and hygiene, 32(5), 1108–1119.

14. Peinado, S. A., Aliota, M. T., Blitvich, B. J., & Bartholomay, L. C. (2022). Biology and Transmission Dynamics of Aedes flavivirus. Journal of Medical Entomology.

15. McLean, B. J., Hall-Mendelin, S., Webb, C. E., Bielefeldt-Ohmann, H., Ritchie, S. A., Hobson-Peters, J., … & van den Hurk, A. F. (2021). The insect-specific Parramatta River virus is vertically transmitted by Aedes vigilax mosquitoes and suppresses replication of pathogenic flaviviruses in vitro. Vector-Borne and Zoonotic Diseases, 21(3), 208–215.

16. Contreras-Gutierrez, M. A., Guzman, H., Thangamani, S., Vasilakis, N., & Tesh, R. B. (2017). Experimental infection with and maintenance of cell fusing agent virus (Flavivirus) in Aedes aegypti. The American journal of tropical medicine and hygiene, 97(1), 299.

17. Bolling, B. G., Olea-Popelka, F. J., Eisen, L., Moore, C. G., & Blair, C. D. (2012). Transmission dynamics of an insect-specific flavivirus in a naturally infected Culex pipiens laboratory colony and effects of co-infection on vector competence for West Nile virus. Virology, 427(2), 90–97.

18. Saiyasombat, R., Bolling, B. G., Brault, A. C., Bartholomay, L. C., & Blitvich, B. J. (2011). Evidence of efficient transovarial transmission of Culex flavivirus by Culex pipiens (Diptera: Culicidae). Journal of medical entomology, 48(5), 1031–1038.

19. Lutomiah, J. J., Mwandawiro, C., Magambo, J., & Sang, R. C. (2007). Infection and vertical transmission of Kamiti river virus in laboratory bred Aedes aegypti mosquitoes. Journal of Insect Science, 7(1).

20. Baidaliuk, A., Lequime, S., Moltini-Conclois, I., Dabo, S., Dickson, L. B., Prot, M., … & Lambrechts, L. (2020). Novel genome sequences of cell-fusing agent virus allow comparison of virus phylogeny with the genetic structure of Aedes aegypti populations. Virus evolution, 6(1), veaa018.

21. Martin, E., Tang, W., Briggs, C., Hopson, H., Juarez, J. G., Garcia-Luna, S. M., … & Hamer, G. L. (2020). Cell fusing agent virus (Flavivirus) infection in Aedes aegypti in Texas: seasonality, comparison by trap type, and individual viral loads. Archives of Virology, 165(8), 1769–1776.

22. Bolling, B. G., Vasilakis, N., Guzman, H., Widen, S. G., Wood, T. G., Popov, V. L., … & Tesh, R. B. (2015). Insect-specific viruses detected in laboratory mosquito colonies and their potential implications for experiments evaluating arbovirus vector competence. The American journal of tropical medicine and hygiene, 92(2), 422.

23. Stollar, V., & Thomas, V. L. (1975). An agent in the Aedes aegypti cell line (Peleg) which causes fusion of Aedes albopictus cells. Virology, 64(2), 367–377.

24. Patterson, E. I., Kautz, T. F., Contreras-Gutierrez, M. A., Guzman, H., Tesh, R. B., Hughes, G. L., & Forrester, N. L. (2021). Negeviruses reduce replication of alphaviruses during coinfection. Journal of Virology, 95(14), e00433–21.

25. Patterson, E. I., Villinger, J., Muthoni, J. N., Dobel-Ober, L., & Hughes, G. L. (2020). Exploiting insect-specific viruses as a novel strategy to control vector-borne disease. Current opinion in insect science, 39, 50–56.

26. Gu, J. B., Dong, Y. Q., Peng, H. J., & Chen, X. G. (2010). A recombinant AeDNA containing the insect-specific toxin, BmK IT1, displayed an increasing pathogenicity on Aedes albopictus. The American journal of tropical medicine and hygiene, 83(3), 614.

27. Ren, X., Hoiczyk, E., & Rasgon, J. L. (2008). Viral paratransgenesis in the malaria vector Anopheles gambiae. PLoS pathogens, 4(8), e1000135.

28. Ward, T. W., Jenkins, M. S., Afanasiev, B. N., Edwards, M., Duda, B. A., Suchman, E., … & Carlson, J. O. (2001). Aedes aegypti transducing densovirus pathogenesis and expression in Aedes aegypti and Anopheles gambiae larvae. Insect molecular biology, 10(5), 397–405.

29. Ma, Q., Srivastav, S. P., Gamez, S., Dayama, G., Feitosa-Suntheimer, F., Patterson, E. I., … & Lau, N. C. (2021). A mosquito small RNA genomics resource reveals dynamic evolution and host responses to viruses and transposons. Genome research, 31(3), 512–528.

30. Weger-Lucarelli, J., Rückert, C., Grubaugh, N. D., Misencik, M. J., Armstrong, P. M., Stenglein, M. D., … & Brackney, D. E. (2018). Adventitious viruses persistently infect three commonly used mosquito cell lines. Virology, 521, 175–180.

31. R Core Team R: a language and environment for statistical computing. R foundation for statistical computing, 2020. Available: https://www.r-project.org/

32. Wickham H ggplot2: elegant graphics for data analysis. New York: Springer-Verlag, 2016. https://ggplot2.tidyverse.org

33. Jeffries, C. L., White, M., Wilson, L., Yakob, L., & Walker, T. (2020). Detection of Cell-Fusing Agent virus across ecologically diverse populations of Aedes aegypti on the Caribbean island of Saint Lucia. Wellcome Open Research, 5.

34. Ajamma, Y. U., Onchuru, T. O., Ouso, D. O., Omondi, D., Masiga, D. K., & Villinger, J. (2018). Vertical transmission of naturally occurring Bunyamwera and insect-specific flavivirus infections in mosquitoes from islands and mainland shores of Lakes Victoria and Baringo in Kenya. PLoS Neglected Tropical Diseases, 12(11), e0006949.

35. Natal Fernandes, L., de Moura Coletti, T., Julio Costa Monteiro, F., Octavio da Silva Rego, M., Soares D’Athaide Ribeiro, E., de Oliveira Ribeiro, G., … & Charlys da Costa, A. (2018). A novel highly divergent strain of cell fusing agent virus (CFAV) in mosquitoes from the Brazilian Amazon region. Viruses, 10(12), 666.

36. Yamanaka, A., Thongrungkiat, S., Ramasoota, P., & Konishi, E. (2013). Genetic and evolutionary analysis of cell-fusing agent virus based on Thai strains isolated in 2008 and 2012. Infection, Genetics and Evolution, 19, 188–194.

37. Espinoza-Gómez, F., López-Lemus, A. U., Rodriguez-Sanchez, I. P., Martinez-Fierro, M. L., Newton-Sánchez, O. A., Chávez-Flores, E., & Delgado-Enciso, I. (2011). Detection of sequences from a potentially novel strain of cell fusing agent virus in Mexican Stegomyia (Aedes) aegypti mosquitoes. Archives of virology, 156(7), 1263–1267.

38. Cook, S., Bennett, S. N., Holmes, E. C., De Chesse, R., Moureau, G., & De Lamballerie, X. (2006). Isolation of a new strain of the flavivirus cell fusing agent virus in a natural mosquito population from Puerto Rico. Journal of General Virology, 87(4), 735–748.

39. Werling, K. L., Johnson, R. M., Metz, H. C., & Rasgon, J. L. (2022). Sexual transmission of Anopheles gambiae densovirus (AgDNV) leads to disseminated infection in mated females. bioRxiv.

40. Cross, S. T., Maertens, B. L., Dunham, T. J., Rodgers, C. P., Brehm, A. L., Miller, M. R., … & Stenglein, M. D. (2020). Partitiviruses infecting Drosophila melanogaster and Aedes aegypti exhibit efficient biparental vertical transmission. Journal of Virology, 94(20), e01070–20.

41. Mao, Q., Wu, W., Liao, Z., Li, J., Jia, D., Zhang, X., … & Wei, T. (2019). Viral pathogens hitchhike with insect sperm for paternal transmission. Nature communications, 10(1), 1–10.

